# Convolutional neural network, personalised, closed-loop Brain-Computer Interfaces for multi-way control mode switching in real-time

**DOI:** 10.1101/256701

**Authors:** Pablo Ortega, Cédric Colas, Aldo Faisal

## Abstract

Exoskeletons and robotic devices are for many motor disabled people the only way to interact with their environment. Our lab previously developed a gaze guided assistive robotic system for grasping. It is well known that the same natural task can require different interactions described by different dynamical systems that would require different robotic controllers and their selection by the user in a self paced way. Therefore, we investigated different ways to achieve transitions between multiple states, finding that eye blinks were the most reliable to transition from ‘off’ to ‘control’ modes (binary classification) compared to voice and electromyography. In this paper be expanded on this work by investigating brain signals as sources for control mode switching. We developed a Brain Computer Interface (BCI) that allows users to switch between four control modes in self paced way in real time. Since the system is devised to be used in domestic environments in a user friendly way, we selected non-invasive electroencephalographic (EEG) signals and convolutional neural networks (ConvNets), known by their capability to find the optimal features for a classification task, which we hypothesised would add flexibility to the system in terms of which mental activities the user could perform to control it. We tested our system using the Cybathlon BrainRunners computer game, which represents all the challenges inherent to real time control. Our preliminary results show that an efficient architecture (SmallNet) composed by a convolutional layer, a fully connected layer and a sigmoid classification layer, is able to classify 4 mental activities that the user chose to perform. For his preferred mental activities, we run and validated the system online and retrained the system using online collected EEG data. We achieved 47, 6% accuracy in online operation in the 4-way classification task. In particular we found that models trained with online collected data predicted better the behaviour of the system in real time suggesting, as a side note, that similar (ConvNets based) offline classifying methods present in literature might find a decay in performance when applied online. To the best of our knowledge this is the first time such an architecture is tested in an online operation task. While compared to our previous method relying on blinks with this one we reduced in less than half (1.6 times) the accuracy but increased by 2 the amount of states among which we can transit, bringing the opportunity for finer control of specific subtasks composing natural grasping in a self paced way.

## I. INTRODUCTION

For many severely paralysed users, such as those suffering from Spinal Cord Injury, Multiple Sclerosis, Muscular Dystrophy or Amyotrophic Lateral Sclerosis, exoskeletons and robotic devices often constitute the only way to interact meaningfully with their environment. In addition, these natural interactions are represented by dynamical systems with different properties whose control can be better achieved by, first identifying the task at hand, and then, providing the best controller for it. However the way in which this can be implemented in a natural and user friendly way remains challenging [1]. Our lab has previously developed a 3D eye-tracking based robotic arm able to assist in reaching movements in the 3D space [2], [3]. In this system the binocular gaze point is used to find the reaching target aimed by the user assisted by the robotic arm and a trigger signal is used to switch between ‘off’ and ‘reaching’ modes. Our group previously investigated electromyography (EMG), voice and eye blinking modalities as triggers, finding that blinks approaches focusing on end-users requirements, specifically algorithms that work online responding to user’s intent in short time. The BrainRunners game (figure 1) for the Cybathlon’s BCI-race [5] requires four commands whose decoding accuracy determines the velocity of an avatar facing an equal number of obstacles indicated by a colour, and represents the challenges of any real time4-command control task yet providing an standarised framework for comparison of solutions.

**Fig. 1.**
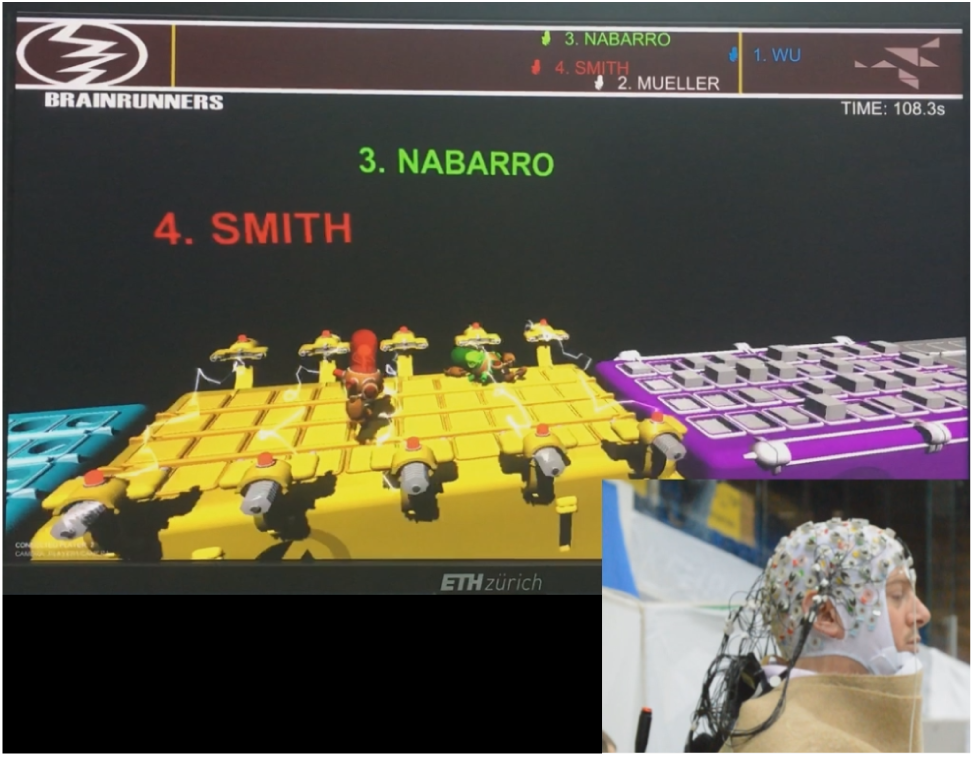
Snapshot from pilot’s point of view of BrainRunners video-game. Right bottom corner shows the EEG set up on our pilot. Each avatar correspond to a user competing in the race. Each obstacle is indicated by a different colour: cyan for avatar rotation; magenta for jump; yellow for slide; and grey for ‘off’ mode. The control is achieved by decoding different mental tasks associated with each desired command. Source: *Cybathlon BCI Race 2016*.

Our system is devised to be used by people with a broad spectrum of disabilities. Therefore, rather than fixing the mental activities and create a classifier that would work for those particular activities and EEG features, we chose Convolutional Neural Networks (ConvNets) to find the optimal personalised EEG features for a user. These architectures are known by their capabilities to extract the best features from offline EEG data for particular mental tasks [6], [7]. We expand on this showing that they are flexible enough to perform online classification of those mental tasks preferred by the user.

It is noted that although several approaches have been proposed for BCI-EEG [8], [9], [10], most of them limit to offline analyses. Furthermore, Deep Learning for EEG signals decoding has also been used in [11], [12] for clinical studies but the architectures provided are computationally demanding and would render real time decoding unsuitable.

Focused in online BCI use constraints we (1) investigated different ConvNet architectures that led us to select a simple one —or SmallNet— made of one convolutional layer, one fully connected layer and a logistic regression classification layer. To overcome the reduced abstraction capabilities of our architecture, we reduced the complexity and size of the network input compared to the raw signal. In particular, we used spectral power features preserving the spatial arrangement of electrodes. Second, (2) we exploited topographical and spectral differences of EEG activities related to 8 different mental tasks that one volunteer found easy to perform and found a combination of four rendering better classification accuracies. Finally, based on our results from previous steps (3) we carried out adaptive training. To the best of our knowledge this is the first time a ConvNet has been tested in an online four-way classification task. In [7] a similar approach is taken but the ConvNet used consists of four layers for a binary classification and there is no online testing of the architecture proposed. Our main contribution consists on the design and implementation of a BCI based on an efficient ConvNet architecture achieving over random accuracies on 4 classes in real use conditions, establishing a baseline for ConvNet based real time BCI implementations within the standarised online control framework of Cybathlon, and bringing the possibility to increase the number of control states a user can switch between in a self paced way only using brain signals.

The remaining sections of this paper are organised in parallel to the three stages of our approach. We start describing the general aspects of data acquisition, and the methods to perform the described analysis. Continuing with the results of our proposed approach. And concluding with the discussion of the results and the limitations.

## II. METHODS

EEG data was recorded using a BrainVision ActiCHamp ^®^ (v. 1.20.0801) recorder. 64 electrodes were placed using the 10-20 system using ‘Fpz’ as reference. Electrooculogram (EOG) activity was recorded on the right eye to correct for ocular artifacts using independent component analysis (ICA) from the *MNE* Python toolbox [15], [16]. ICA matrices were computed offline and the rows corresponding to EOG components were removed from the matrix. This way the reconstruction of the signal using this modified matrix was used to clean the online generated data, ensuring the algorithm was using only brain activities and not ocular artefacts.

The input size to the convolutional layer (CL) depended on the preprocessing method applied to the raw EEG. This preprocessing consisted in the topographical arrangement of the EEG’s power spectrum (Figure 2). 129 frequencies spanning from 0 to 250Hz at 1.95Hz intervals were used. For each frequency an image was created by arranging the corresponding power of each channel to an approximate topographical 2D projected distribution of the channels.

**Fig. 2.**
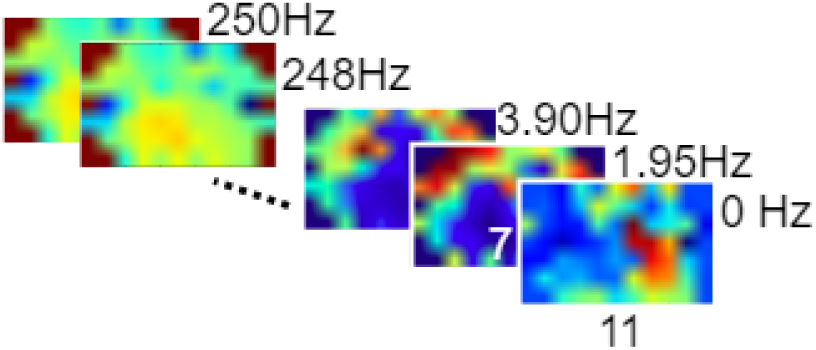
Input features. 129 spectral power images of the 2D EEG channel projection. The tensor input size to the ConvNets resulted in [129,11,7].

The architectures were fed by data the same way during offline training and online decoding. Each 300ms a 1.2s chunk of EEG (with 0.9 already used seconds and 0.3 *fresh* seconds accounting for 75% overlap) was used by the algorithm. In the offline case each of the described EEG chunks configured a training example.

ConvNet architectures were implemented in Theano [13] and the system was built in an Intel i7-6700 CPU at 3.40GHz with model’s training and execution running in a NVIDIA 1080GTX. A 28 year old, right handed man volunteered throughout all the stages and all the experiments were approved by the Imperial College Ethical Committee. Unless explicitly stated significance alpha level is 0.01 for the analysis.

### A. Architecture selection-Stage 1

The tested architectures are presented in figure 3. Different complexities (building up from SmallNet, Figure 3 B) of ConvNets were tested with the intention of finding one able to abstract enough information from our limited set of examples. For each run, the weights were randomly initialised following a uniform distribution within the [-1,1] range.

**Fig. 3.**
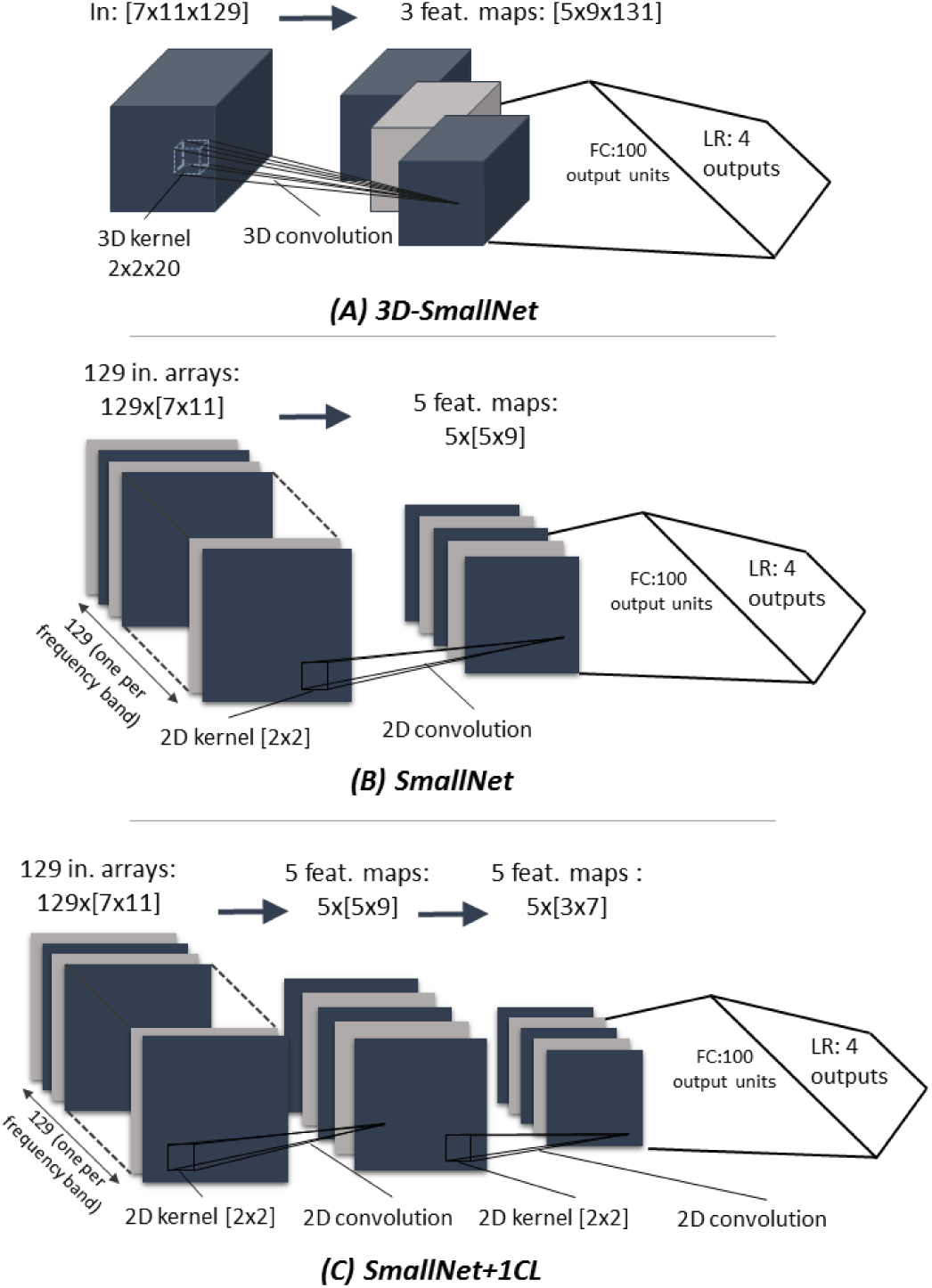
Three different ConvNets architectures: A 3D-SmallNet (A) using 3D convolution instead of the 2D convolution used by SmallNet (B). A convolutional layer was added to Small-Net (C, SmallNet+1CL) and also a fully connected after the first convolutional one not presented in the figure (SmallNet+1FC).

All the architectures were trained using a dataset acquired as follows. The user was instructed to perform contraction imageries corresponding to the feet, stomach, right hand and left hand. He performed these imageries while watching 20 previously recorded videos of the race with a balanced but randomly distributed amount of pads. Whenever the avatar in the video got into a new pad the user had to continuously perform the mental activity assigned to that pad. Examples were extracted as indicated in the Methods introduction. A chronologically arranged 5-fold validation approach was conducted. The selection of the chronological arrangement of folds ensured that examples corresponding to the training and testing sets were not overlapped, leading to overfitting. Around 9000 examples were extracted this way.

### B. Mental tasks evaluation-Stage 2

The best resulting architecture was then used to select a combination of four mental tasks providing best classification results. The combinations were taken from a set of 8 imageries that the user felt comfortable performing, spanning: motor imageries (right hand, left hand, lips or feet contraction), higher cognitive processes (mental humming and arithmetics) and relax. An experimental paradigm, with screened instructions instead of videos, was used to acquire 100 1-second examples per each mental activity and used to train 70 (8-choose-4) models using the best architecture following the same k-fold strategy.

### C. Adaptive training design-Stage 3

The last stage sought to validate and analyse the results in real time conditions and investigate an adaptive online training providing feedback to the user. The importance of this kind of adaptation has been already addressed in [14].

#### Validation stage

To evaluate the online use the following strategy was devised. First, a recording session of 20 videos, similar to that in the previous section, was used in the same manner to train SmallNet (Fig. 4). A second session, immediately after, used that model for online decoding during five races made of 20 pads. Both the race time, and two decoding accuracies were used to analyse the results. One decoding accuracy (*acc*_1_) considered the label corresponding to the pad where the EEG data was generated and the other (*acc*_2_) considered that the decoded label arrived in the correct pad. They could differ if a long decoding time prevented the decoded label for one data portion to arrive on the pad it was generated. The same previous training parameters were used in this stage.

**Fig. 4.**
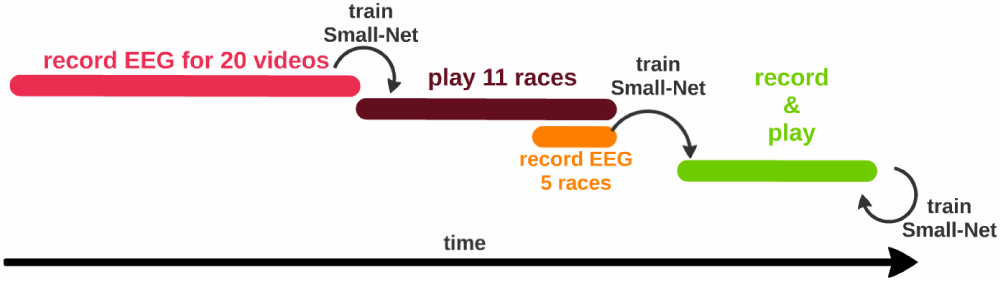
Non-adaptive (warm colours) and adaptive training (green) strategies. At the beginning of the session EEG activities are recorded during 20 videos. These were used to train the SmallNet and this model was used to play for 11 races. EEG data in playing conditions from the last 5 races were recorded and used to retrain the model to start the adaptive training. After each adaptive training race data was recorded, appended to those 5 races and the model retrained.

#### Adaptive training

An old model was used to decode the commands for the first race, meanwhile the data generated was stored and used to train and validate another model for posterior races. This second model was updated after each race appending to previous examples the new data generated. A limit of 2000 was used to train the model dropping data from the oldest race each time a newer one was appended to limit the time the model spent training between races.

## III. RESULTS

### A. Architecture selection-Stage 1

A first analysis discarded 3D-SmallNet due to its long training time, making it impractical for an adaptive online approach. For the rest of the architectures, the 5-fold strategy was run 5 times per architecture to control for the effect of different initialisations on accuracy (Table I). Because there was no clear benefits of adding more layers, we used SmallNet as it required the shortest training time.

**Table I.** TEST ERROR (TE) OF RELEVANT ARCHITECTURES

| | $mt_4$ | $TE_{avg}$ | $TE_{std}$ |
| --- | --- | --- | --- |
| 1 | SmallNet | 58.21 | 1.34 |
| 2 | SmallNet+1CL | 53.52 | 8.56 |
| 3 | SmallNet+1FC | 58.89 | 1.92 |

### B. Mental tasks evaluation-Stage 2

A Kruskall-Wallis test revealed significant differences between groups of mental tasks combinations. Table II shows the number of combination groups that each of the presented combination shows pairwise significant differences with.

**Table II.** PAIRWISE COMPARISONS OF IMAGERIES COMBINATIONS

|  | im1 | im2 | im3 | im4 | # sig. diff. |
| --- | --- | --- | --- | --- | --- |
| 1 | RH | feet | lips | numbers | 18 |
| 2 | RH | feet | relax | numbers | 16 |
| 3 | RH | feet | relax | lips | 16 |
| 4 | RH | relax | lips | stomach | 11 |
| 5 | RH | feet | stomach | numbers | 11 |
| 6 | RH | feet | music | numbers | 10 |
| 7 | RH | relax | stomach | numbers | 10 |
| 8 | RH | relax | lips | numbers | 10 |
| 9 | LH | feet | relax | lips | 8 |
| 10 | RH | LH | relax | lips | 6 |
| 18 | LH | feet | relax | stomach | 3 |
| 19 | RH | feet | relax | m. humming | 3 |

To respect the user friendly design aim and given that there was no best combination in absolute terms (i.e. no combination group was significantly better than all the rest) we let the volunteer choose the one which he felt more comfortable with when playing: **RH-feet-relax-music**. It still presents some advantages over 3 groups and is not statistically different from any combination above it (*TA*_*mean*_ = 46.85% for the selected group compared to 55.01% for group 1 in Table II).

### C. Adaptive training-Stage 3

Figure 5 shows the results for the online session. *Acc*_1_, and *acc*_2_ are the validation accuracies measured as explained in methods, while *acc*_*test*_ is the offline test accuracy of the model used to decode during playing conditions (ref. to Figure 4 for training strategy).

**Fig. 5.**
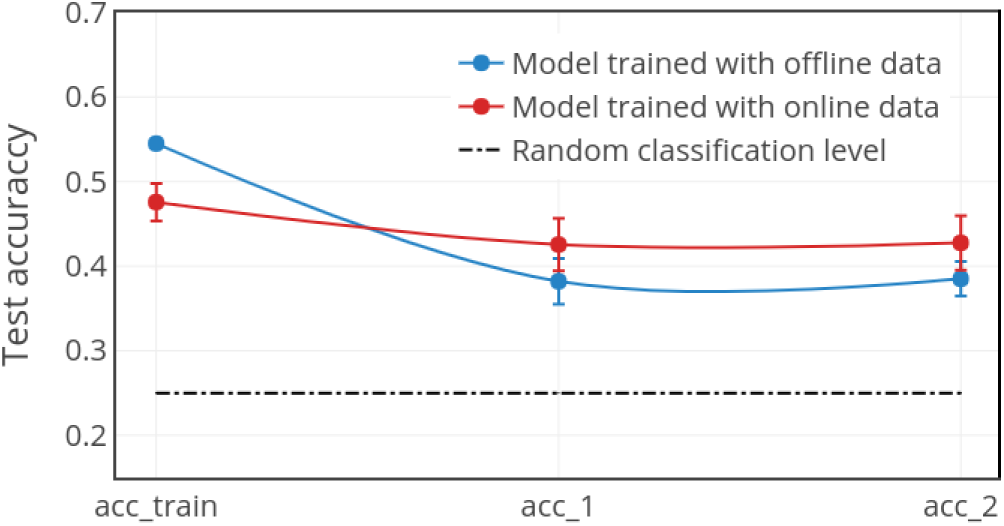
Online playing test accuracies. Playing results for the model trained with offline data (non-adaptive) are presented in red. Those on blue correspond to the model trained with data gathered in actual playing conditions (adaptive).

Several tests were carried out analysing aspects related to the online function. First, no differences existed between the two methods to measure accuracies: *acc*_1_ and *acc*_2_.

In terms of accuracy there is no differences between the adaptive and the non-adaptive training. Indeed both accuracies yielded by both strategies (adaptive and non-adaptive) correlate similarly to the time invested on each race. Their Pearson’s coefficients (accuracy, time) being, -0.385 and -0.388 respectively, thus both carrying a similar reduction in time when accuracy is increased (Figure 6).

**Fig. 6.**
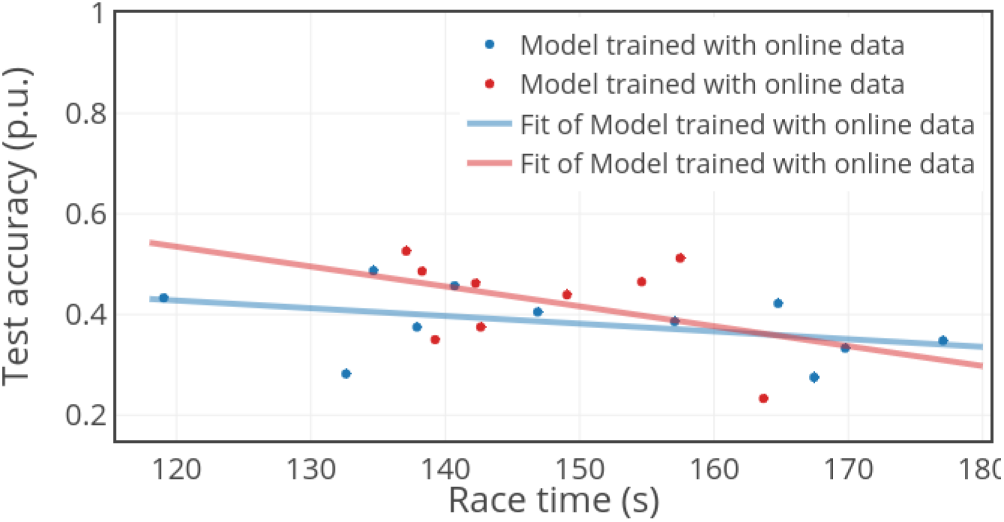
Time VS validation accuracy. The figure shows the results in decoding accuracy (*acc*_2_) during the game and the time required to finish the race. A linear fit is applied to show the correlation between both values.

However, the online training predicted the online behaviour much better than the offline one. While test accuracies for adaptive training (*acc*_*test*_ = 0.476) were not significantly different to those achieved online, the accuracies achieved during offline training were systematically higher than those achieved in using the offline trained model in real time (*p* = 7.14·10^*-5*^). This is a crucial aspect defending the real time validation of such systems, supporting the idea that commonly offline reported results often overestimate the achievable real time performance.

## IV. DISCUSSION

### A. Architecture selection-Stage 1

We have showed the capabilities of an efficient ConvNet (SmallNet) to distinguish among four different brain activities in a single example basis, achieving significant over random accuracies (*>* 25%). We tested variations departing from a very simple ConvNet architecture in order to find if the increase in computing complexity could led to higher accuracies. For our particular setup and dataset we found that SmallNet, being the simplest model, provided similar accuracies than more complex ones justifying its use.

Training ConvNets depends very much on the initialisation when data is scarce. We run several times the same training in order to get estimates of the variability and found that the impact is important to consider. Methods to better initialise the architecture should be further investigated to achieve consistent convergences.

### B. Mental tasks evaluation and features-Stage 2

The state of mind can change from in the interval of hours. We collected data for the study of mental tasks days before the online validation. In addition, because we wanted our subject to feel comfortable, we let him choose the combination of mental tasks that he felt more comfortable with, however it remains to clarify if other combinations would have presented different results.

A previous study of input features evidenced that spectral power features were, regardless of the data fold and the initialisation of the network weights, better than any of the *raw-time* features. Spectral energy features in channel space have been classically used to characterise and study brain activities in several frequency bands, as they have been found to enhance statistical differences. In addition given the high sampling frequency, for the same amount of time, the computation of such features greatly reduced the input dimension, which enable the small architecture to learn more efficiently. However the space to explore features is enormous and affect different aspects of the system, like training time and consistency and decoding accuracy. We would sought for features that lead to shorter training times and reduce the time that the user needs to invest prior to the use of the system. In our case we finally considered that our choice was good given its simplicity and low computational demands, but others should be further studied.

### C. Adaptive training validation-Stage 3

We validated the system using the volunteer preferred imageries (*RH-feet-relax-music* in the top-19). In particular we found that adaptive test accuracies offered a more reliable prediction of validation accuracies during playing. A reasonable explanation for this is that adaptive training uses EEG brain activity collected in playing conditions, the same that are present when the model is used to decode. Conversely, training the model with videos EEG recordings yielded better test accuracies in data sets with more examples but less representative of the brain activity during actual playing. We demonstrated that for a ConvNet based BCI, adaptive training can achieve same performance as offline training while engaging the participant since the beginning.

To this respect, one of our main concerns is the relevance of the ICA matrix along time. For online playing we computed it only once at the beginning. Therefore we consider that how good this ICA matrix is at correcting eye artifacts may change along time and not be as representative for future eye activity sources as time passes. A possible solution would be to recompute it after each race in the adaptive training just before training the model. However, differences in convergences of the matrices across races may introduce some differential distortions in the back projected data that hinder the ability of the network to effectively learn. Finally, as more races are used in the adaptive training more examples are available for the net to learn and more time the training takes. Shorter set-ups are important for domestic and user friendly implementations. We consider relevant to study the effect of limiting the number of examples to train the network to stablish a limit in training time after each race. Another concern is the stability of the model along races. Although overall performance is good from race to race there is variation in accuracies across races. We think it is important to understand how models can be transferred from one race to another and find if the variations are caused by user changes in focus rather than model or recording instabilities. Finally and as previously mentioned, our major concern relates to how offline results can be translated to online ones. Our offline analyses gave us some clues about what architecture and mental task to use but testing all combination online would be impractical which supports the idea of letting the user choose among the top performer combinations. Our online results for 4 classes are comparable to those reported in [20] where only half the participants achieved around 65% of accuracy in a more time-locked experiment compared to our user-driven real timeset. In [21] using a real time binary BCI the drop in online accuracy compared to offline was *∼*16% in average compared to 29.9% for our non-adaptive training and 8.8% for adaptive which situate in the margins of the first figure, showing that the drop in accuracy is somewhat generalisable.

## V. CONCLUSION

In a context where a consistent and standard benchmark for testing is lacking we have presented a rational approach to the design and implementation of a BCI fulfilling the real time control requirements. We identified two usual weaknesses in BCI and DL as decoding technique: (1) in most BCI conceptions users performance evoking a richer range of brain signals is only summarily considered; and (2) in general most research involving DL focus on offline evaluations of decoding accuracy, leading to complex architectures whose impact in real life conditions is rarely discussed. We took these improvement opportunities to design an experimental protocol based on a ConvNets selected by its simplicity and the low computing resources it required so that it could be used in real time and would not require long setup times. In particular, we showed that more complex architectures did not provide any advantage in terms of accuracy which may suggest that our approach is limited by the number of examples and thus more complex architectures do not offer an advantage.

Our results confirmed the value of considering different brain activities categories in order to increase accuracy. We also found evidence suggesting that preprocessing methods rendering high frequency-resolution topographical-energy features of brain activity improve the capacity of our small architecture to distinguish among categories.

We found that the adaptive training strategy yielded a more realistic representation of actual playing accuracies, though the accuracy achieved (*∼*55%) is still far from what users need. the exploitation of different imageries and features is a reasonable source of accuracy improvement.

In conclusion, by using this noninvasive approach we were able to double the number of states our robotic arm could be in (representing different control modes) but nearly halved the accuracy when comparing to the blinking method we previously used. These preliminary results show the potential of SmallNet to increase the number control states completely independent from muscular activity, broadening the spectrum of motor disabilities such a system could be applied to. Further experiments are to be carried to see how the system apply to a broader range of users.

## ACKNOWLEDGMENT

We thank BrainProducts for the loan of the 160 channel ActiCHamp EEG recorder. We also appreciate and thank the organisation of Cybathlon 2016. The support of the EPSRC Centre for Doctoral Training in High Performance Embedded and Distributed Systems (HiPEDS, Grant Reference EP/L016796/1) is gratefully acknowledged during the last stages of this work.

